# A probabilistic model-based bi-clustering method for single-cell transcriptomic data analysis

**DOI:** 10.1101/181362

**Authors:** Sha Cao, Tao Sheng, Xin Chen, Qin Ma, Chi Zhang

## Abstract

We present here novel computational techniques for tackling four problems related to analyses of single-cell RNA-Seq data: (1) a mixture model for coping with multiple cell types in a cell population; (2) a truncated model for handling the unquantifiable errors caused by large numbers of zeros or low-expression values; (3) a bi-clustering technique for detection of sub-populations of cells sharing common expression patterns among subsets of genes; and (4) detection of small cell sub-populations with distinct expression patterns. Through case studies, we demonstrated that these techniques can derive high-resolution information from single-cell data that are not feasible using existing techniques.

## Introduction

Single cell sequencing represents a new generation of sequencing technique, allowing the capture of genomic and transcriptomic differences in a cell population, consisting of possibly different cell types. Compared to the currently available RNA-Seq data for tissue samples such as those in the TCGA database [1], which measure the averaged expression levels of genes across all the cells, single-cell RNA-Seq data contain gene-expression levels of individual cells, hence enabling studies of behaviors of individual cells or cell types as well as their complex interactions under specific conditions. Using cancer as an example, such new data will enable studies of a wide range of issues that are currently infeasible such as (1) how stromal and some immune cells co-evolve with the diseased cells under persistent stressful conditions such as oxidative stress or acidosis; (2) what specific conditions each key cancer-related mutation such as TP53 or Ras mutation is selected in specific tissues to overcome, hence gaining detailed understanding of how selected mutations contribute to the development of a cancer; and (3) what specific conditions, as reflected by the expression levels of some genes in some cell types, drive a cancer to metastasize.

The single-cell sequencing technique has been applied to study various challenging biological problems in the past few years, such as studies of stem cell differentiation trajectories [2], embryonic development [3-7], distinctive cellular responses to external stimuli [8], lineage hierarchy construction in a whole tissue [9], intra-tissue heterogeneity [10-12], and cancer metastasis origin using circulating tumor cells (CTCs) [13], all of which are feasible only because of the single cell sequencing technique.

Along with these new exciting possibilities also come new challenges in effectively analyzing and interpreting single-cell data. We have identified a number of challenges based on our own experience in using such data, which, we believe, are of fundamental importance and practical generality. Here we present our developed novel techniques to address four such problems. The first challenge is that a target cell population may consist of multiple cell types, suggesting the high possibility of genes showing different behaviors in different cell types, hence the need for a mixture model of multiple distributions for each gene [14, 15]. Some work has been done on this particular issue. For example, Shalek et.al modeled single cell RNA-Seq data by using a bimodal distribution, in which an expression threshold is applied to distinguish samples that significantly express a gene from those that do not [8]. This over-simplistic approach has a number of intrinsic issues such as a gene’s expression pattern could be of more than two peaks, which has limited its applications. The second challenge lies in the unquantifiable errors in the large number of observed zeros or low expression values in a single-cell expression profile, causing traditional Gaussian models not directly applicable. A third challenge was to detect sub-populations of cells sharing common expression patterns by some (to be identified) subsets of genes, which may represent different subtypes of cells that have not been previously characterized. For example, our previous study has discovered that cancer tissues in general consist of two distinct cancer cell types, one being more anaerobic and the other being more like normal (aerobic) cells [16]. Our own experience has been that detection of sub-populations of cells sharing common expression patterns of some genes represents a widely encountered problem, and in general it can be modeled as a bi-clustering problem. A fourth challenge lies in detecting relatively small cell sub-populations with significantly distinct expression patterns. Principle component analysis (PCA) has been a popular approach to tackling this problem [17, 18]. However the basic assumption needed for using PCA, namely the orthogonality assumption may not be accurate since all cell types in a tissue are generally not orthogonal in their expressions; instead they are likely to be related.

In this paper, we present a mixed Gaussian distribution with left truncated assumption, named LTMG, to model the single cell RNA-Seq data to address the first two challenges. To the best of our knowledge, this model is the first rigorous statistical model to fit gene expression profiles with a large number of zeros/low expression data that can accurately capture the expression patterns of a gene through multiple single cells. We also developed a procedure for converting a mixed Gaussian distribution into a binary, 1/0, string, representing genes being differentially expressed or not, on a scientifically sound basis for bi-clustering analysis using our in-house bi-clustering tool QUBIC [19]. This novel bi-clustering procedure proves to be especially effective in coping with time-and/or location-dependent single-cell data. The mixed Gaussian distribution also makes it apparent whether a cell is overly expressing a specific gene, which provides a basis for identification of small cell population bearing rare biological characteristics. We have implemented these novel techniques as a computer analysis pipeline, and demonstrated the effectiveness of these techniques and the pipeline through conducting three case studies.

## Results

### Analysis pipeline for transcriptomic data

In this study, we developed a novel distribution to fit single-cell gene-expression profile under multiple conditions, along with an analysis pipeline *LTMG-QUBIC*, based on the distribution, for identifying significantly expressed genes in individual cells, clusters of cells with similar expression profiles over a (not pre-defined) subset of genes, and outlier cells showing high expressions in a distinctively large number of genes.

The analysis pipeline consists of four steps, as illustrated in Figure 1: (I) log-transformed RPKM (Reads Per Kbp of transcripts) [20] values of each gene measured from single cell RNA-Seq experiment is first fitted by a Left Truncated Mixed Gaussian (LTMG) distribution; (II) the fitted model is then applied to evaluate significantly expressed genes and outlier cells; (III) the fitted distributions are then used to identify statistically significant bi-clusters in a specified dataset using our in-house bi-clustering tool QUBIC; and (IV) functional enrichment analysis is applied to infer the functional characteristics of individual cells and bi-clusters. More details of the analysis pipeline are listed as follows.

**Figure 1.**
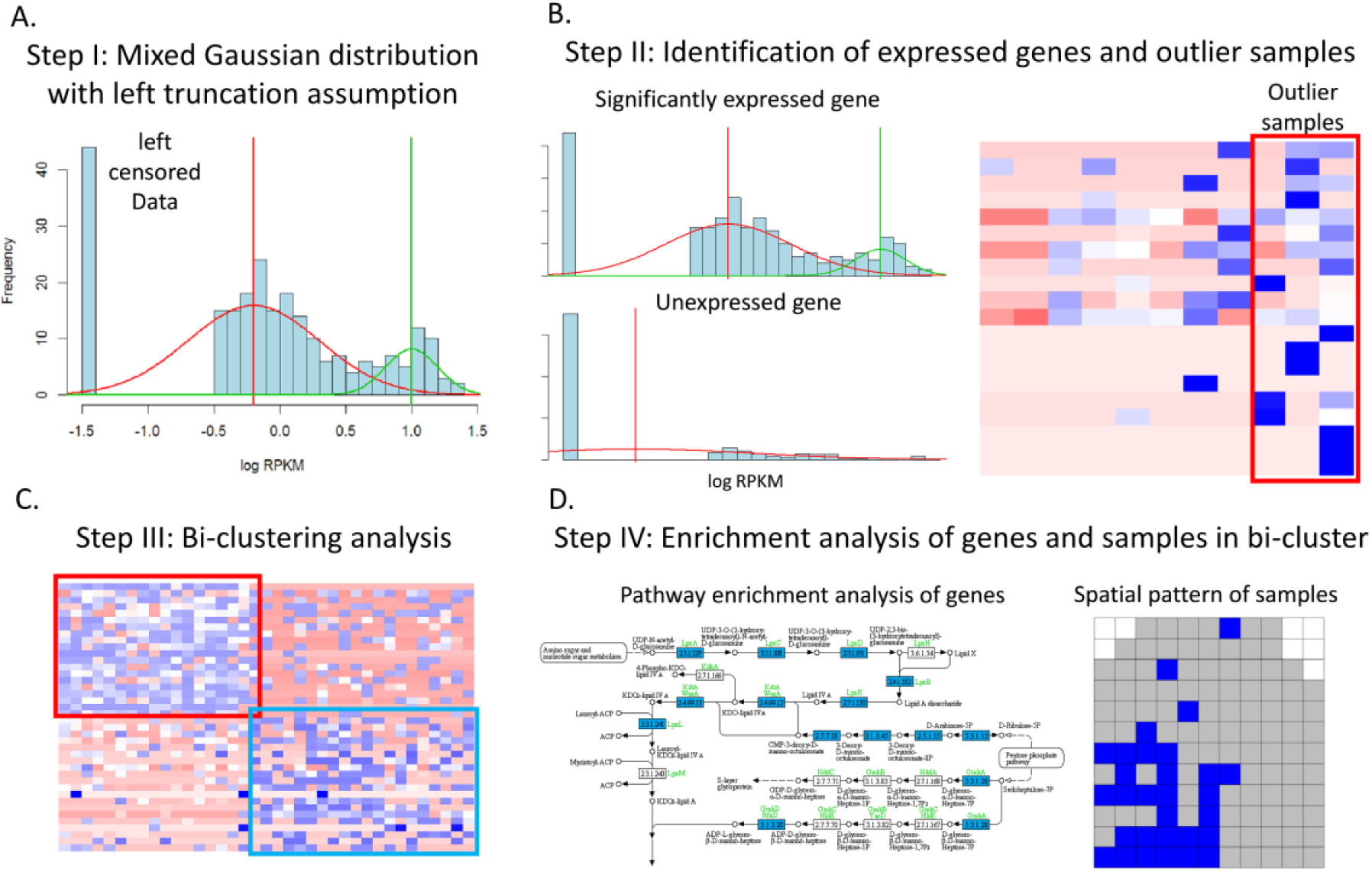
The analysis pipeline. (A) Left truncated mixed Gaussian distribution to fit the log-transformed RPKM single-cell gene expression profile. In the histogram of the log-transformed RPKM gene expression data, the left most bar shows the number of left censored data while the red and green curves represent the two Gaussian distributions identified by using our LTMG model. (B) The left panels show the gene expression profile of one significantly expressed gene (upper) and one non-expressed gene (lower) with estimated mean value smaller than threshold Zcut; the right panel shows a heatmap of the gene expression data of outlier and non-outlier samples, in which low and high expressions are represented by red and blue colors, respectively. (C) Identified significant bi-clusters using QUBIC, where the heatmap shows the gene expression data of two identified bi-clusters. (D) Pathway enrichment analysis is applied to capture high-level functions of each identified bi-cluster. The left panel shows genes significantly enriched in certain pathways and the right presents the spatial pattern of the samples in one bi-cluster identified in the spatial transcriptomic data.

### Step I: Mixed Gaussian model with left truncation assumption to model single cell RNA-Seq profile

Single cell RNA-Seq samples collected under different conditions may give rise to an expression distribution with more than one peak for each gene [14, 15]. Hence a mixed distribution needs to be employed to model an expression profile across multiple cell types, especially for genes that only express in some subsets of the RNA-Seq samples [21]. Noting the large number of zero values in single cell RNA-Seq data may cause unquantifiable errors in fitting the distribution to the expression profile, we employed a left truncation assumption to model the log-transformed RPKM expression values by mixed Gaussian distributions [22]. Specifically, the expression values lower than a threshold are treated as left censored data when the gene expression profile is fitted using a mixed Gaussian distribution as shown in Figure 1A. The detailed information regarding how a gene-expression profile is fitted by a mixed Gaussian distribution with left truncation assumption using the EM algorithm is given in the Material and Methods section.

### Step II: Identification of significantly expressed genes and outlier samples

One challenging problem in single cell RNA-Seq data analysis is to determine if a gene is truly expressed when small PRKM values are observed. Using our LTMG model, a gene is considered as *significantly expressed* when the mean of the fitted distribution is greater than a preselected threshold when the distribution has only one peak, or the distribution has more than one peak. More details about the gene-specific threshold for significant expression is given in the Material and Methods section. The non-expressed genes were excluded from further analyses.

Another challenging issue in single cell RNA-Seq data analyses is to reliably identify outlier cells defined by cells with a large number of highly expressed genes, and such samples may represent certain rare events such as stem cells or early response to stimuli in the immune system as in our case studies. Since the expression profile of each gene has its mixed Gaussian distribution, thus for any individual sample, we can assess whether its expression of the gene is significantly high by computing the probability of obtaining an expression value equal or greater than what was actually observed for the sample based on the fitted distribution. By doing this, we can count the number of genes with significantly high expression in each sample, and an outlier sample is called if it has significantly large number of such highly expressed genes, and the significance of such an observation can be evaluated by using permutation tests. (See the Material and Methods section for details).

### Step III: Bi-clustering analysis to identify significant co-expression patterns

It has been well recognized that bi-clustering analyses are essential to discovery of novel biological pathways and functionally associated genes, which offer more powerful analysis capabilities than the traditional one-dimensional clustering techniques [23]. The key idea is to find subsets of all genes and subsets of all cells that exhibit common expression patterns. Here we convert each gene-expression level to a binary number, 0 or 1, representing no or differential expression as required by our bi-clustering software QUBIC, based on a rule detailed in the Material and Methods section. Then a bi-cluster is defined as an all “1” sub-matrix of the 0/1 matrix representing the expression levels of genes (rows) in specific cells or under specific conditions (columns).

### Step IV: Linking bi-clusters to cell type or spatial specific expression patterns

Genes and conditions in each identified bi-cluster are then analyzed by pathway enrichment analyses to infer higher-level functional information associated with specific cells or cell groups.

### Application of the analysis pipeline on single cell and spatial transcriptomic data

We applied our analysis pipeline on three recently published datasets: (1) GSE48968, a single-cell RNA-Seq time-course dataset of 1,700 primary mouse dendritic cells treated with three pathogenic agents and multiple control experiments [24]; (2) GSE57872, a single cell RNA-Seq dataset of 430 glioblastoma cells isolated from five distinct tumors with known clinical information [12]; and (3) GSE60402, an bRNA-Seq dataset of laser-capture micro-dissected tissue cubes from the medial ganglionic eminence of wild-type and GFRa1 mutant mice where each cube containing ∼100 cells was treated as a single cell with detailed spatial coordinates [25]. More detailed information on the three datasets could be found in the Material and Methods section. Each of the three datasets contains multiple sub-datasets collected under different conditions as summarized in Table 1 with more detailed information given in Supplementary Table S1-S3. The numbers of significantly expressed genes, outlier samples and bi-clusters were identified using the analysis pipeline on each sub-dataset.

**Table 1.** The numbers of significantly expressed genes, genes with multiple distribution peaks, outlier samples, and bi-clusters in each analyzed data set.

| **Data ID** | **#significantly expressed genes** | **#genes with multiple distribution peaks** | **#significant outlier samples** | **#significant bi-clusters** |
| --- | --- | --- | --- | --- |
| GSE60402-Mutant | 8,589 | 8,497 | 24 | 37 |
| GSE60402-Wildtype1 | 6,957 | 6,887 | 12 | 40 |
| GSE60402-Wildtype2 | 8,600 | 8,490 | 15 | 120 |
| GSE57872-MGH26 | 4,490 | 3,823 | 7 | 44 |
| GSE57872-MGH28 | 5,426 | 4,768 | 17 | 131 |
| GSE57872-MGH29 | 5,558 | 4,339 | 16 | 43 |
| GSE57872-MGH30 | 5,425 | 4,698 | 16 | 17 |
| GSE57872-MGH31 | 5,146 | 4,062 | 12 | 42 |
| GSE57872-ALL | 5,814 | 5,584 | 109 | 460 |
| GSE48968-LPS | 9,980 | 9,872 | 123 | 51 |
| GSE48968-PAM | 8,451 | 8,307 | 71 | 24 |
| GSE48968-PIC | 8,834 | 8,697 | 90 | 17 |

### GSE48968

We first used our LTMG model to analyze the single cell time-course RNA-Seq data of the 1,700 primary mouse dendritic cells (DCs) collected at 1h, 2h, 4h and 6h after treatment with three pathogenic agents, namely LPS (lipopolysaccharide), PAM (synthetic mimic of bacterial lipopeptides) and PIC (viral-like double-stranded RNA) and untreated controls [24]. On average, 9,088 genes were found to be significantly expressed, 94 outlier cells and 31 bi-clusters are identified in the three pathogen-treated datasets. One question raised in the original work is to identify the genes with significant bimodal distributions. By our LTMG model, 52.21% (7306/13991) genes were fitted as bimodal and 18.34% (2566/13991) genes are fitted as multimodal distribution in the data of LPS treatment while the percentages are 52.95% (6951/13127) and 10.32% (1356/13127) in the PAM data and 52.46% (7152/13632) and 11.33% (1545/13632) in the PIC data, respectively (see Table 1). This shows the advantage of our method assuming a variable number of peaks for single cell gene expressions compared with the original study.

Principle component analysis was applied in the original study to identify time-dependent immune responses by assuming a fixed number of time-dependent patterns. However, the number of true time-dependent patterns could be much larger than the estimated fixed number. By applying our LTMG-QUBIC analysis, all the possible time-dependent immune responses were comprehensively identified by bi-clustering analyses that simultaneously cluster genes and cells sharing a common time-dependent pattern. In total, 51, 24 and 17 statistically significant bi-clusters were identified in the datasets treated with LPS, PAM and PIC, respectively. For each bi-cluster, the Fisher exact test was conducted on its constituting samples to assess if significant over-representation by any time points could be found within the bi-cluster. For those bi-clusters showing significant association with the time-course, a pathway enrichment analysis on genes of the bi-cluster is conducted to infer the biological characteristics of the bi-cluster. At the end, 30 bi-clusters that are significantly over-represented by one or several consecutive time points have been identified in the LPS dataset using *α*=0.005 as the significance cutoff and six of them with distinct time dependence (p<1e-22) are presented in Figure 2.

Specifically, bi-cluster BC013 consists of untreated samples and samples collected at 1h after LPS treatment, which represents the earliest response to LPS, and enriches multiple immune response pathways such as T-cell activation and interleukin signaling pathway (Figure 2A). Bi-cluster BC005 consists largely of untreated samples and samples collected at 1h and 2h after the LPS treatment, which also enrich immune response pathways but more responses to virus, T cell chemotaxis, NK cell activation, and regulation of NF-kB (Figure 2B), representing further responses compared to BC013. The immune response pathways enriched in these two bi-clusters suggest that the immune response is the earliest response to the LPS treatment in dendritic cells. BC009 (Figure 2C) and BC001 (Figure 2D) are enriched by samples collected at 1h and 2h after the LPS treatment, which include a wider range of stress-response pathways such as hemostasis, apoptotic mitochondrial changes, glucose metabolism, lipid and fatty acid metabolism, NOTCH, P53 and HIF signaling and immunological synapse formation pathways, suggesting that the activation of stress response pathways and altered metabolisms as secondary responses after the early immune response. BC025 (Figure 2E) is enriched with samples collected at 4h and 6h after the LPS treatment, whose genes enrich pathways of NOTCH transcription and translation, NIK/NF-kB signaling, regulation of cell migration, endothelial cell morphogenesis, hematopoietic progenitor cell differentiation, and regulation of stress-activated MAPK cascade. Another bi-cluster BC002 consists of largely samples collected at 4h and 6h after the LPS treatment with genes that enrich pathways related to growth factor binding, lipopolysaccharide-mediated signaling, cell membrane, cell junction, cell-cell adhesion, leukocyte chemotaxis, and calcium ion homeostasis pathways (Figure 2F). In addition, both BC025 and BC002 genes enrich pathways associated with alterations in cell morphogenesis, migration and cell-cell junction. Overall these observations suggest that our analysis is capable of identifying all the major responses to the LPS treatment in a time-dependent manner. Detailed pathways enriched by the six bi-clusters are given in Figure 2.

Similarly, time-dependent immune responses are identified from 11 and 13 time-course associated bi-clusters in the PAM and PIC datasets. The detailed information of these bi-clusters is given in Supplementary Figure S1 and Supplementary Table S1.

Another key question addressed in the original study of this dataset is to identify cells showing early response to the pathogens at each time point. As a comparison, we pooled the samples at two adjacent time points (unstimulated and 1h, 1h and 2h, 2h and 4h, and 4h and 6h) with the same pathogenic agent treatment, and applied the analysis pipeline on four pooled datasets. In total, three, three, seven and one significant outlier cells are identified in the datasets collected at 1h, 2h, 4h and 6h after the LPS treatment, respectively. Pathways enriched by the significantly expressed genes in the outlier cell samples include cell’s response to interleukin and interferon signaling, ion homeostasis and cell-cell junction, suggesting that these outlier cells represent stronger response to the LPS treatment comparing to the other cells at the same time interval. Similarly, outlier cell samples responding to PAM and PIC treatments are also identified. Specifically, two outlier cells show early immune response with highly expressed immune responsive pathways among the 2h cells in the PAM data and 1h cells in the PIC data, which are consistent with the reported results in the original study. The detailed information of the identified outlier samples is given in Supplementary Table S1.

**Figure 2.**
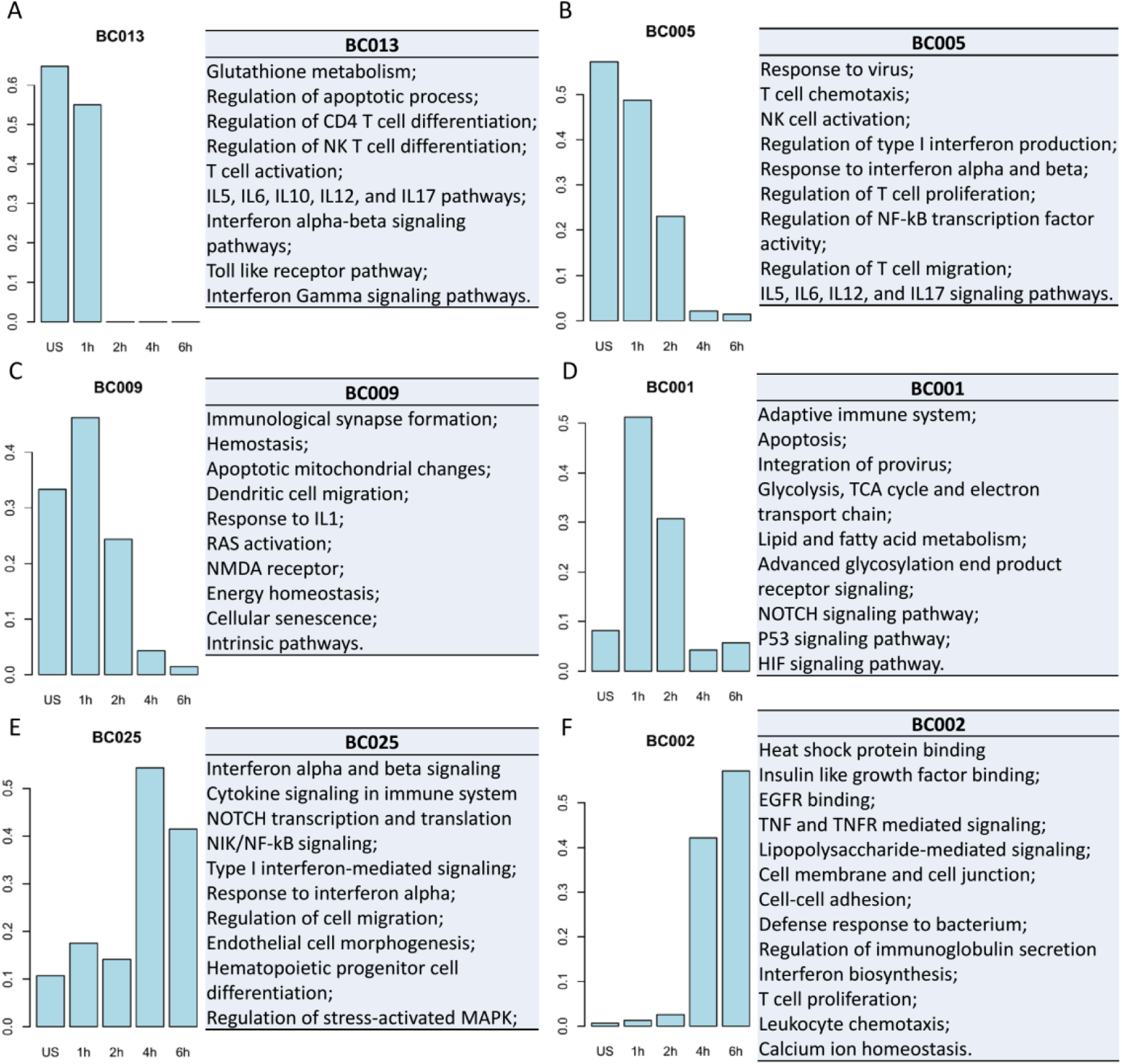
Time-dependent distribution of samples and enriched pathways by genes in six selected bi-clusters identified in the LPS data. In each panel, the five bars from left to right show the proportion of the untreated samples and samples collected at 1h, 2h, 4h and 6h after the LPS treatment.

### GSE57872

We then applied our pipeline on the dataset of 430 single-cell samples collected from five glioblastoma tumors to allow a systematic characterization of intra-tumor and inter-tumor variability as well as identification of stem-like cells. Specifically, our bi-clustering analysis of the data set of each tumor captures the intra-tumor variability associated gene co-expression patterns while inter-tumor variability associated co-expression patterns were identified by using the data from all five tumors. Outlier identification method was then applied to target stem-like cells. On average, 87.58%(5,209/5,948) genes are identified as significantly expressed gene, 14 cells are predicted as outliers and 35 bi-clusters are identified in the dataset of each tumor while 97% (5,814/5,948) significantly expressed genes with 109 outliers and 679 bi-clusters are identified in the merged data set (Table 1). It is noteworthy that only 5,948 genes were measured in the original study.

The most frequently observed pathways in the above bi-clusters are related to cell cycle phases, glucose, fatty acid, amino acid and glycan metabolisms and hypoxia response genes, suggesting that the intra-tumor variability is majorly reflected on the cell proliferation and metabolism level, which is consistent with the original report of the dataset [12]. In addition, our method identified numbers of novel pathways showing significant intra-tumor variability in each tumor including proteasome and NF-kB activation in MGH26, cellular export machinery, N-linked glycosylation and post translational modification in MGH28, mRNA decay, cellular export machinery and steroid synthesis in MGH 29, cholesterol biosynthesis, cellular export machinery and MYC pathway in MGH30, and proteasome, amino acid metabolism and regulation of apoptosis in MGH31.

460 statistically significant bi-clusters were identified in the pooled dataset, in which 413 were significantly enriched by samples from one or several tumors (tested by the Fisher exact test with *p*-value < 0.005) that are considered as inter-tumor variability associated gene expression patterns. Pathway enrichment analyses on these bi-clusters revealed that the most frequently observed inter-tumor variability associated pathways cover TP53 signaling, EGFR signaling, MTOR and AMPK signaling, glycosaminoglycan metabolism, cell cycle, lipids metabolism, cell adhesion, apoptosis, immune response and central metabolism pathways. Specifically, glycolysis and TP53 signaling pathways are identified in one bi-cluster corresponding to tumor MGH29 with TP53 mutation. EGFR signaling, G2-M phase in cell cycle and lipid metabolism pathways are identified in bi-clusters composed of samples from MGH26 and MGH31, both of which harbor copy number variations of EGFR. Cells from MGH28 and MGH29 with wild type EGFR show distinct expression patterns in the pathways of immune response, CD8 T cell and MHC class 1 complex, compared with the rest of the tumors in the dataset. Bi-clusters corresponding to the samples in MGH26, MGH29, MGH28, MGH28_29_30, MGH26_31, MGH26_28_31, MGH29_30, MGH28_29, and MGH26_30_31 with enriched pathways are selected and presented in Figure 3.

We identified approximately 14 outlier cells in the data of each tumor. In each of the five tumors, 4-8 outlier cells are enriched by highly expressed cell cycle genes and at least one cell with highly expressed stem cell marker genes. In addition, the most frequently observed pathways that are enriched by the highly expressed genes in outlier cells include: innate immune response and interferon gamma signaling, TCA cycle and electron respiration chain, cell cycle and stem cell marker.

**Figure 3.**
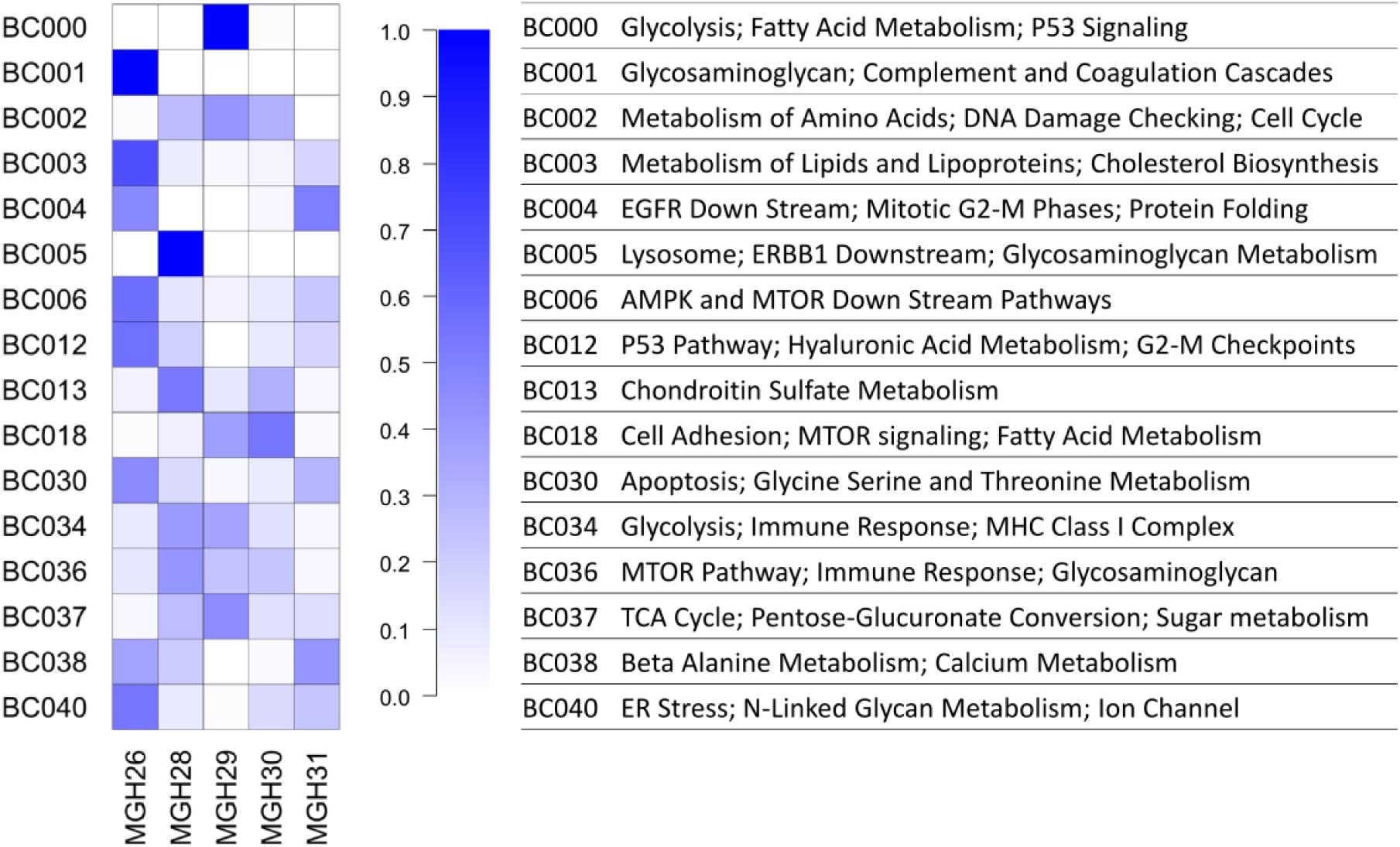
Enriched pathways of genes in selected bi-clusters identified in GSE57872 dataset, and the tumor tissues that the samples in these bi-clusters come from. The heatmap represents the proportion of the samples in each tumor identified in each bi-cluster with color key given in the right.

### GSE60402

This dataset contains samples dissected from three mouse medial ganglionic eminence tissues with known spatial coordinates that enable the analysis of location-dependent characteristics such as cell differentiation trajectories in neuron development. Each biological sample in the dataset contains approximately 100 cells, which were treated as a single cell when generating their transcriptomic data (see Material and Methods). Our bi-clustering and outlier analysis were applied on the dataset to identify location-dependent cell clusters and spatial distribution of cells showing unique expressions of neuron development related genes. Our analysis identified 75.67% (8,048 out of 10,037) genes as significantly expressed, 17 outlier cells and 136 bi-clusters out of the three datasets corresponding to the three different mice (See Table 1 for details).

37, 40, and 120 bi-clusters were identified in the mutant, wild type 1 and wild type 2 datasets, respectively, and analyzed regarding the spatial distribution of cells in each bi-cluster. All the four spatial clusters with distinct expression patterns by cell cycle, cell morphogenesis and neuron development genes, as reported in the original study, were identified by our bi-clustering and pathway-enrichment analysis (Figure 3A, 3C) [25]. In addition, our method discovered more distinctions across the four clusters as follows: cytokine-cytokine receptor, mitochondrial fatty acid beta oxidation, phospholipid metabolism, telomere maintenance, galactose metabolism, histidine metabolism, O-linked glycosylation, ribosome, spliceosome, and circadian.

In addition to the four major clusters, more than 50% of the identified bi-clusters tend to contain samples from nearby spatial locations. Our analysis has identified 19, 23 and 48 additional bi-clusters with samples covering cells in adjacent spatial regions in the three data sets as shown in Supplementary Figure S2. Pathway enrichment analysis revealed that these bi-clusters mostly enrich pathways of TCA cycle, respiratory electron transport, oxidative phosphorylation, signaling by WNT, Huntington’s disease, Alzheimer’s disease, Parkinson’s disease, regulation of apoptosis, chromatin modification, cell-cell adhesion, and translation, suggesting spatial dependent variabilities of these pathways.

12, 15 and 24 significant outlier samples were identified in the mutant, wildtype 1 and wildtype 2 data, respectively. Pathway enrichment analysis revealed that most of these outliers have highly expressed genes that enrich the cellular functions such as histone H3 tri-methylation marker at K27, high-CpG-density promoters (HCP) with un-methylated histone H3, stem cell markers, cell cycle, oligodendrocyte development and differentiation, suggesting a possible association between stem properties and specific methylation pattern of H3. It is noteworthy that the outliers with highly expressed stem cell markers tend to be located at the intermediate region between two adjacent (or overlapping) bi-clusters in the three datasets as shown in Figure 3B and 3D. Our interpretation is that these location-dependent expression patterns may be caused by parallel and independent differentiations from common stem cells.

**Figure 4.**
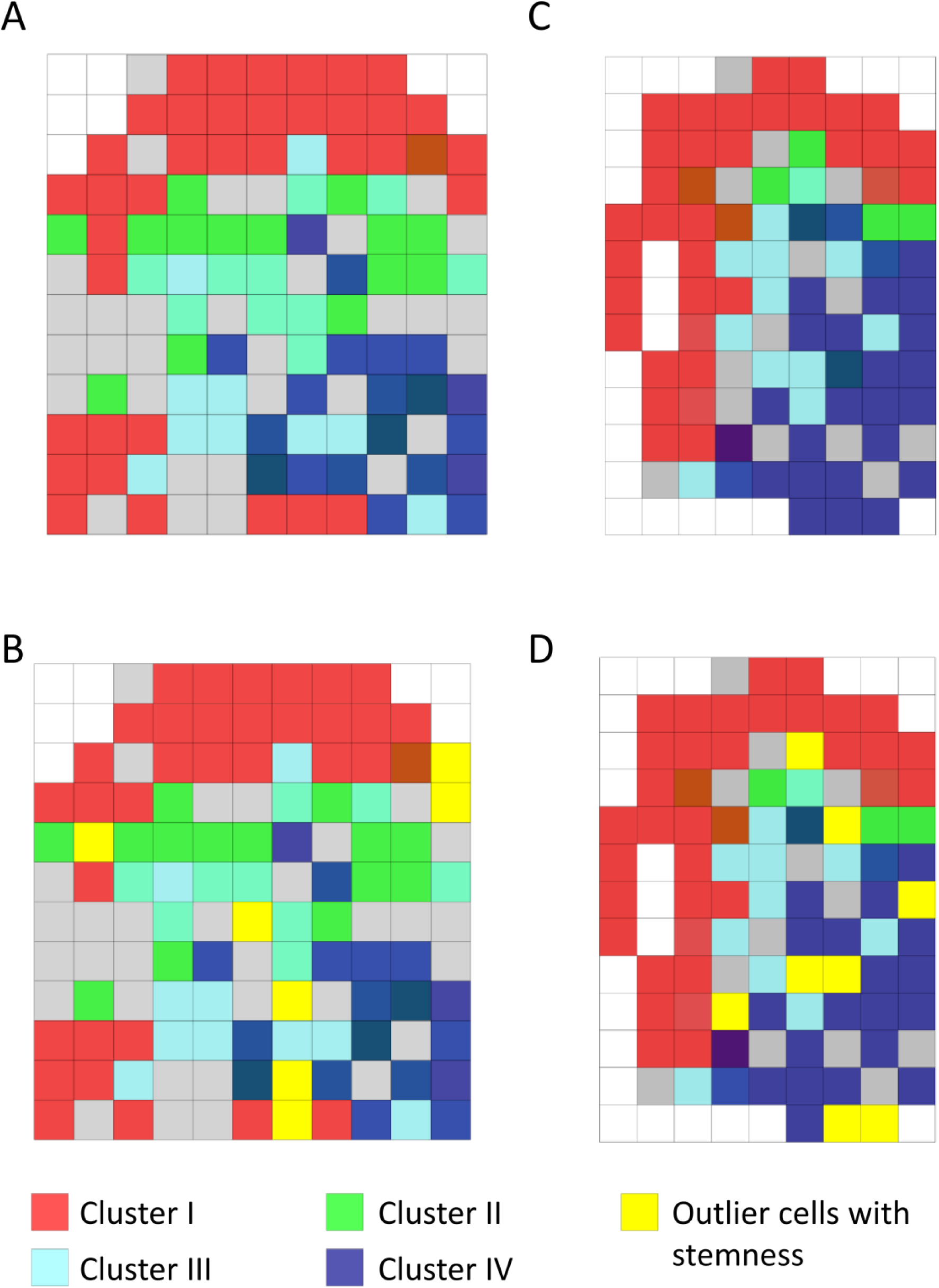
The spatial coordinates of samples in identified bi-clusters and outlier samples in the wild type 1 (A and B) and wild type 2 (C and D) datasets. A) The spatial coordinates of samples in the four bi-clusters identified in wild type 1 single cell samples. Colors red, green, cyan and dark blue represent samples in four different bi-clusters. B) In addition to the coordinates of bi-cluster samples, the yellow cubes represent significant outlier samples. C) The same information as in A) except the samples are from wild type 2 mouse. D) The same information as in D) except the samples are from wild type 2 mouse.

## Conclusion

A computational analysis pipeline for single-cell RNA-Seq data has been developed and presented here. The pipeline consists of four novel methods for tackling challenging analysis problems of single-cell RNA-Seq data, including (1) effective modeling of multimodal geneexpression profiles across multiple cell types; (2) handling the unquantifiable errors due to the large number of zeros and low-expression levels often observed in single-cell RNA-Seq data; (3) reliable and efficient identification of sub-populations of cells sharing common expression patterns by some not pre-selected genes; and (4) detection of relatively small cell sub-populations with significantly distinct gene expression patterns. We fully anticipate that these techniques and the analysis pipeline will prove to be a useful resource to the analysts and users of single-cell RNA-Seq data in their tissue-or other mixed cell population based studies.

## Discussion

Single-cell sequencing has enabled new transcriptome-based studies, including study of distinct responses by different cell types in the same population when encountered by the same stimuli or stresses, and identification of the complex relationships among different cells in complex biological environments such as tissues. In this paper, we presented a novel computational method for analyses of single-cell based transcriptomic data, which allows to reliably address the following types of questions: (1) which genes in which cells show responses to specific stimuli, through reliable detection of differentially expressed genes in specific cells or cell types; (2) how a cell sub-population responds to specific stimuli, in terms of the common responses by the whole cell sub-population, detected through bi-clustering analyses, as well as outlier responses by small subsets of the sub-population; and (3) location and time-dependent cellular responses. To address these issues reliably, we have to overcome a number of technical challenges.

A key challenge lies in that a target cell population may consist of multiple cell types, suggesting the high possibility of genes showing different behaviors in different cell types. Our mixed Gaussian distribution assumes variable expression patterns for each gene, and effectively captured as much expression patterns as possible of through multiple single cells.

A second challenge arises when the large number of observed zeros or low expression values causes unquantifiable errors in a single cell expression profile, making traditional Gaussian models not directly applicable. This issue is handled through left truncating a Gaussian distribution and treating these truncated values as missing data. To the best of our knowledge, our left truncated mixed Gaussian model is the first rigorous statistical model to fit gene expression profiles with a large number of zeros/low expression data, which can accurately capture the expression patterns of a gene across different cell types

A third challenge, in our view, is to address the need for answering: how different (not pre-determined) sub-populations of cells respond differently to specific stimuli. In essence, this is to solve a bi-clustering problem. We have previously developed a computational program QUBIC, now widely used for solving a bi-clustering problem. In the current study, we developed a procedure for converting a mixed Gaussian distribution into a binary, 0/1, string, to represent no or differentially expressed genes as required by the QUBIC program for bi-clustering analyses. Notably, we extended the bi-clustering analyses to include a new capability to capture co-expressed genes in a time-or location-dependent manner. Our case studies show that the integration of our model with the QUBIC bi-clustering method works well with single cell RNA data under multiple treatment conditions and/or in a temporal/spatial dependence manner.

A fourth challenge is to detect relatively small cell sub-populations with significantly distinct gene expression patterns. Our mixed Gaussian distribution allows for rigorously assessing the statistical significance in observing high expression of a gene in a sample.

By integrating these novel capabilities, we developed a computational pipeline for handling single-cell RNA-Seq data in an automated manner, hence allowing application of our new analysis tools to large quantities of single-cell data through web-based server techniques, which is currently under construction, aiming to serve a large user community.

Through three case studies, we demonstrated that (1) our new method can not only detect what the original study of each of the three datasets detected, but also offer new information not provided by the original studies; (2) our novel distribution for modeling single-cell gene expression profiles provides a statistically rigorous model for single-cell gene expression data across different cell types, and offers an effective framework for identification of outlier cells bearing rare biological characteristics; (3) our bi-clustering method proves to be capable of handling expression data with complex experimental conditions, including both temporal or spatial information.

Through utilizing time-dependent data, our analysis has identified a cascade of immune responses to the external pathogenic treatment. On the glioblastoma data, our analysis detected stem-cell related functionalities as well as intra-tumor variabilities represented by cell proliferation and central metabolism and distinct inter-tumor variability as reflected by glycosaminoglycan metabolism and cell growth signals. On the spatial transcriptomic data, we discovered spatially adjacent single cells may have high co-expression patterns, and particularly, two distinct spatially clustered cells may be originally derived from the same stem cell.

## Material and Methods

### Data selected in the analysis

Datasets GSE57872 and GSE58968 are downloaded from the GEO database and raw reads of GSE60402 are retrieved from SRA database [26, 27]. The RPKM values for the first two datasets are available from the GEO database, and the RPKM values for GSE60402 were calculated by using software packages TopHat [28] and Cufflink [29]. All these single-cell datasets have large sample sizes, and are collected using the latest experimental protocols. Each dataset is split into multiple subsets based on experimental conditions, each of which was analyzed separately. The detailed information of the selected and split datasets are listed in Table 3.

**Table 3.** Data information.

| GEO<br>Accession<br>ID | Data ID | Description | Number<br>of<br>samples | Number<br>of genes |
| --- | --- | --- | --- | --- |
| GSE60402 | GSE60402-Mutant | From Gfra1 mutant sample | 94 | 11094 |
| GSE60402 | GSE60402-Wildtype1 | From wild type mouse 1 | 124 | 10037 |
| GSE60402 | GSE60402-Wildtype2 | From wild type mouse 2 | 94 | 10714 |
| GSE57872 | GSE57872-MGH26 | From Tumor sample MGH26 | 53 | 5948 |
| GSE57872 | GSE57872-MGH28 | From Tumor sample MGH28 | 94 | 5948 |
| GSE57872 | GSE57872-MGH29 | From Tumor sample MGH29 | 75 | 5948 |
| GSE57872 | GSE57872-MGH30 | From Tumor sample MGH30 | 73 | 5948 |
| GSE57872 | GSE57872-MGH31 | From Tumor sample MGH31 | 70 | 5948 |
| GSE57872 | GSE57872-ALL | Complete GSE57872 data set | 430 | 5948 |
| GSE48968 | GSE48968-LPS | LPS treated cell with control | 527 | 13991 |
| GSE48968 | GSE48968-PAM | PAM treated cell with control | 325 | 13127 |
| GSE48968 | GSE48968-PIC | PIC treated cell with control | 419 | 13632 |

### Log-mixed Gaussian model with left truncation assumption to model the expression profile of each gene measured using the RPKM normalized expression level

To accurately model the gene expression profile of single cell data, we specifically developed a mixed Gaussian model with left truncation assumption to fit the log transformed gene expression measured by RPKM. Mixed Gaussian distribution has been widely applied to model multiple peaks in gene expression data, and the model fits very well the single-cell transcriptomic data where a large number of genes express only in some subsets of all the samples in each dataset [30]. Noting that the error of the RPKM expression level at zero or the low-expression values does not fit a normal distribution, we have therefore introduced a left truncation assumption when fitting the mixed Gaussian model [22].

Denoting the observed log-transformed RPKM expression level of gene X over N cells as *X* = (*x*_1_, *x*_2_,…, *x*_*N*_). We assume that *x ∈ X* follows a mixture Gaussian distribution with K components corresponding to K possible peaks and the density function of X is:

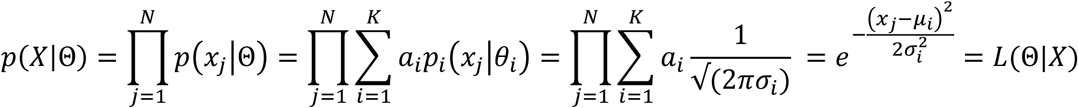

where parameters Θ = {*a*_*i*_, *u*_*i*_ σ_*i*_ | *i* = 1… *K*} and *a*_*i*_, *u*_*i*_ *and σ*_*i*_ are the proportion, mean and standard deviation of each Gaussian distribution, respectively, which can be estimated by the EM algorithm with given datasets X:

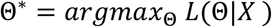

To model the errors at zero and the low expression values, we introduce a parameter Z_cut_ for each gene expression profile and consider the expression values smaller than Z_cut_ as left censored data [22]. With the left truncation assumption, the gene expression profile are split into M truly measured expression value (> Z_cut_) and N-M left censored gene expressions (≤ Z_cut_) for the N conditions. Latent variables *y*_*i*_ and Z_j_ are introduced to estimate Θ by the following Q function and using the EM algorithm:

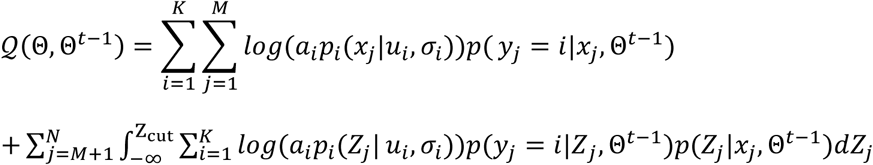
where Θ = {*a*_*i*_, *u*_*i*_ σ_*i*_ | *i = 1…K*} are the parameters, *t* is the current iteration step, Z_cut_ is the cutoff of the measured gene expression level of X to have reliable Gaussian errors; *x*_*j*_ is the measured gene expression level of X, i.e., log-transformed RPKM value in cell j; Z_j_ is the latent variable reflecting the real expression level of X if the measured expression level is smaller than Z_cut_ and *y*_*j*_ is the latent variable reflecting that *x*_*j*_ is the from the *y*_*j*_ th Gaussian distribution.

To estimate the parameters Θ that maximizes the likelihood function, we have Maximization step of the EM algorithm as:

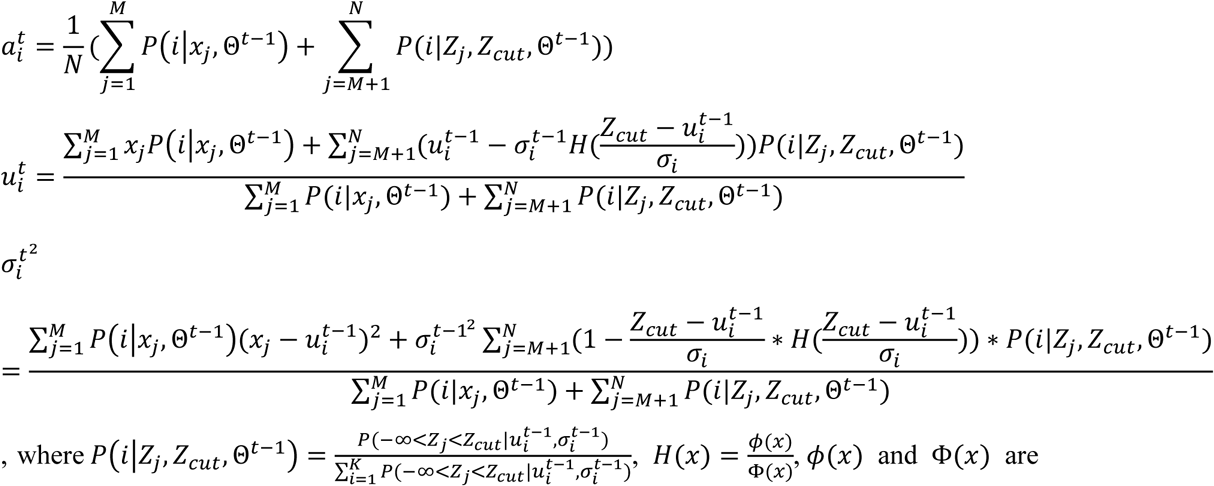

the pdf and cdf of standard normal distributions.

Parameters Θ can be estimated by iteratively running the estimation (E) and maximization (M) steps. The complete algorithm is given in the Supplementary Method. In this study, Z_cut_ is set for each gene as logarithm of the minimal non-zero RPKM value in the gene’s expression profile. The EM algorithm is conducted for K = 1,…, 9 to fit the expression profile of each gene and the K that gives the best fit is selected according to the Bayesian Information Criterion (BIC) [30].

### Assessment of the significantly expressed genes and identification of outliers

For the gene expression profile fit by one Gaussian distribution, the gene is considered as *non-expressed* if the mean value of the distribution is smaller than Z_cut_ while the gene is considered as *expressed* if the mean value is larger than Z_cut_. Note that for the gene expressions fit by multiple Gaussian distributions, they have at least one component with mean value larger than Z_cut_, hence considered as significantly expressed genes. Significantly expressed genes are applied in outlier identification and bi-clustering analysis.

The likelihood of each expression value x_j_ with respect to distribution *i* can be computed by:

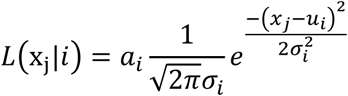

And x_j_ is determined to correspond to component i if 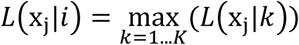

An expression outlier is identified as a sample whose expression is significantly high with respect to the fitted distribution, and the significance, *p*_*j*_, is defined as the probability of observing an expression with a larger value than the currently observed expression *x*_*j*_

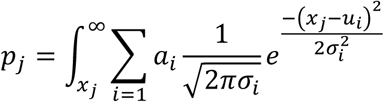

A gene’s expression level is deemed as significantly high in sample *j* if *p*_*j*_< 0.005. A 1,000-time permutation test is performed to find samples with a significantly large number of outlier expressions (called *outlier samples*) and *p*=0.005 is applied as the cutoff for determining an outlier sample. More details of the permutation test are available in the Supplementary Method.

### Data discretization and bi-clustering analysis

To apply our in-house bi-clustering analysis tool QUBIC, the expression profile of each gene is discretized into 1/0 values as required by QUBIC. Specifically, for each gene whose expression profile can be fitted with one Gaussian distribution (with left truncation), the top 10% expressed (non-zero) samples are assigned with 1 and the rest 0. For a gene whose expressions are fitted by a mixture of K (K > 1) Gaussian distributions, K discretized vectors are generated. Then, each sample *i* is assigned to its most probable component calculated as arg 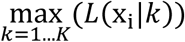 for the gene, and for the *k*th vector corresponding to the gene, its *i*th element takes value 1 only when the *i*th sample’s expression of the gene is from the *k*th component of the fitted mixed Gaussian distribution, and 0 otherwise.

We applied our QUBIC to identify bi-clusters from the discretized data. To ensure high identification specificity, we set the parameter f as 0.25 to ensure that the bi-clusters will not overlap with each other substantially (with other parameters as default). The significance of each identified bi-cluster is estimated by running the bi-clustering procedure on 1000 randomly shuffled discretized data. The S score in the QUBIC output is applied to estimate the *p* value of each bi-cluster; and *p*=0.005 is applied as the significance cutoff. See details of this part in the Supplementary Method.

### Pathway enrichment analysis

Pathway enrichment analysis is conducted and the statistical significance of each enriched pathway is assessed by using a hypergeometric test (statistical significance cutoff = 0.005) against 4,725 curated gene sets in the MsigDB database, which includes 1,330 canonical KEGG, Biocarta and Reactome pathways, and 3,395 gene sets representing expression signatures derived from experiments with genetic and chemical perturbations, together with 6,215 Mouse GO terms each containing at least 5 genes [31, 32].

## Authors’ contributions

CZ, QM and YX designed the statistical model. CZ, SC and XC developed the algorithm and analysis pipeline, and performed the analysis. CZ, SC, QM and YX wrote the manuscript. QM and YX supervised the whole project. All authors read and approved the final manuscript.

